# Ultrastructural preservation of a whole large mammal brain with a protocol compatible with human physician-assisted death

**DOI:** 10.64898/2026.03.04.709724

**Authors:** Aurelia Song, Anna LaVergne, Borys Wróbel

## Abstract

Building a high-fidelity computational model of the whole human brain will require preservation of the ultrastructure at the level of the entire organ, post-mortem. For such a model to reflect as closely as possible the brain in the living state, artifacts that arise during both the agonal phase and the postmortem interval will need to be minimized. This is potentially feasible if a terminally-ill patient donates their brain for research following physician-assisted death. In this paper, we modify a protocol for aldehyde-stabilized cryopreservation to make it compatible with physician-assisted death. We use pigs as a model, which resemble humans in cardiovascular and brain anatomy. Aldehyde-stabilized cryopreservation was designed to provide superior structural preservation of brains of any size, across all anatomical scales, compatible with diverse analytical assays and long-term storage without ultrastructural degradation. We demonstrate, with light microscopy and volume electron microscopy, that our brain preservation protocol results in connectomically traceable whole brains and propose an economically feasible storage modality that is expected to maintain stability of ultrastructure and macromolecules in the brain even for thousands of years. Most importantly, we establish that 14 min is the approximate length of the perfusability window—the time after the cardiac arrest during which blood washout needs to be initiated so that the brain ultrastructure is preserved.

## Introduction

The human brain is hierarchically organized across spatial scales from nanometers (macromolecules, their complexes), through micrometers (synapses, cells), and millimeters (neuronal circuits), to decimeters (whole-brain networks) [1]. It contains an enormous number of cells and synapses—estimates suggest an average of 60–100 billion neurons (and a similar number of glial cells), with the total number of synapses in hundreds of trillions [2,3]. Understanding how the brain generates internal representations, controls movement, stores and retrieves memories, and supports cognition and consciousness will require bridging all these spatial scales, as these properties emerge at the level of the whole organ and, arguably, above. Ultimately, this will mean building high-fidelity computational models—whole human brain/body emulations—models that result in behavior characteristic to a specific individual, in an appropriate environment, because they implement the internal causal dynamics, replicating biological information processing relevant for cognitive processes in a specific brain [4–6].

Current methods to interrogate brain structure at the nanoscale are both destructive and require brains that are preserved postmortem. For data obtained in this manner to reflect the brain as close to the living state as possible, a preservation protocol needs to minimize artifacts that arise during both the agonal phase and the postmortem interval [7]. This is potentially feasible if a terminally-ill patient donates their brain for research following physician-assisted death (PAD)—in particular, after medical aid-in-dying, in which a terminally-ill patient self-administers physician-prescribed lethal medication [8].

In this paper, we modify a protocol for aldehyde-stabilized cryopreservation (ASC, [9]) of a large mammalian brain to make it compatible with PAD. Because these brains are perfused with aldehydes, ASC allows for ultrastructural preservation of whole brains of any size, and a wide range of methods of structural neuroscience can then be used downstream. Additionally, because cryoprotectant is perfused after initial fixation, these brains can be stored long-term at low temperatures without degradation of ultrastructure or macromolecules [9].

We validate this modified ASC protocol using pigs as a model organism. Pigs closely resemble humans in cardiovascular anatomy [10]. The structure of the heart and the aorta are similar; similar body mass yields comparable hemodynamic parameters and perfusion flow rates. Despite their smaller brain size, pig brains have neuroanatomical features relevant to human brain preservation—the brain is gyrencephalic and maintains a white-to-gray matter ratio similar to humans [10]. The cerebral blood flow in both species is similar: 60 ml/min/100 g in pigs [11] and 50 ml/min/100 g in humans [12]. For these reasons, pigs are commonly used as a model in cerebrovascular research, including studies of ischemic stroke [13].

This issue of ischemia is highly relevant for human brain preservation after PAD, because this protocol necessarily will involve some ischemic time, even if all efforts are taken to minimize it. Such ischemic time is expected to elicit processes similar to those involved in the no-reflow phenomenon after ischemic stroke—the clinical manifestation of microcirculation disturbance that occurs within minutes after the cessation of oxygenated blood flow. These processes include microthrombus formation, edema, pericyte constriction, and stimulated vasoconstriction [14,15]. The cellular and molecular mechanisms responsible for these processes are expected to be similar in pigs and humans, whose evolutionary lines diverged 80–90 Mya [16]. The main result of this paper, therefore, is that the length of the *perfusability window*—the time after the cardiac arrest during which blood washout needs to be initiated so that the brain ultrastructure is preserved—is about 14 min.

## Materials and Methods

### Brain preservation

The brain preservation procedure followed a modified protocol for ASC [10]. All animal handling was in full compliance with USDA standards and guidelines for animal care. Briefly, female Yorkshire pigs (55–70 kg, n=5) were sedated with 5 mg/kg each of telazol, xylazine, and ketamine and then injected with 250 IU/kg of heparin. After ten minutes to ensure adequate heparin circulation, the animals were injected with 3120-3900 mg pentobarbital sodium and 400-500 mg phenytoin sodium. Immediately following cardiac arrest, which happened in less than 1 min after this injection, the heart was exposed; a cannula was introduced into the ascending aorta, rapidly secured, and de-aired; and the flow of washout solution was initiated.

To assess whether this surgical procedure would be feasible with humans, we used a human cadaver donated for scientific research. To enable comparison to the pig experiments, we measured the time from the start of surgery to the end of cannula de-airing, which is when perfusion would have begun. Assessment of brain ultrastructure in this case was irrelevant due to the long pre-existing post-mortem interval.

All percentages in the descriptions of solutions in this section are weight per weight (w/w). After washout (1–10 min; with 1.3 % Na_2_HPO_4_, 0.23 % NaH_2_PO_4_, 0.10 % NaN_3_) and open-circuit fixation (10–30 min; with 3.3 % glutaraldehyde, 1.3 % Na_2_HPO_4_, 0.23 % NaH_2_PO_4_, 0.10 % NaN_3_, 0.010 % SDS) removed most of the blood, the output was recirculated back into the pig, and fixation continued. In two experiments (pigs D and E) this was followed by slow addition of cryoprotectant-fixative solution (with 65.5 % ethylene glycol, 3.0 % glutaraldehyde, 1.2 % Na_2_HPO_4_, 0.21 % NaH_2_PO_4_, 0.091 % NaN_3_). To keep the total volume of the perfusate constant in the closed circuit, part of the effluent was continuously removed at the same rate as that at which the cryoprotectant-fixative solution was added. The presence of NaN_3_ prevents mitochondrial swelling [9,17]. The addition of SDS to the fixative solution prevents excessive osmotic shrinking of the brain during the subsequent addition of cryoprotectant [9], observed also by Rosene et al. when cryoprotectant was perfused after fixation [18]. This is most likely because SDS increases the blood brain barrier permeability [19].

### Tissue processing; sample preparation for electron microscopy

Brain sections were sent to outside laboratories (American Histology, Meyer Instruments) for paraffin embedding, hematoxylin and eosin staining, and scanning.

Samples for electron microscopy (150 μm slices) were stained following a modified Knott et al. protocol [20]. Cryoprotectant, if present, was removed, and then samples were washed three times (2 min each) with the washout solution, and then stained with 1.5% potassium ferrocyanide with 1% osmium tetroxide (30 min), and then with 1% osmium tetroxide (30 min) in the washout solution. Then, they were rinsed twice with distilled water (2 min), stained with 1% uranium acetate (30 min) in the dark, and again rinsed with distilled water (2 min).

The samples were dehydrated in 50%, 70%, 95%, and then three times with 100% ethanol (2 min each); and again three times with propylene oxide (3 min each). After that, they were embedded, first using a 1:1 mixture of propylene oxide with Spurr resin (30 min), then 1:2 such mixture (30 min), and finally, 100% Spurr resin overnight. Small sections were cut from the slices, placed in bullet molds with fresh Spurr resin, cured in an oven at 60°C for at least 48h, and sent to an outside laboratory (Janelia Research) for imaging of serial 50 nm sections collected on a silicon wafer with scanning electron microscopy (wafer-SEM), and focused ion-beam scanning electron microscopy (FIB-SEM).

## Results

In our preliminary experiments, we used four pigs to optimize the perfusion circuit and the surgical technique. These experiments demonstrated that rapid surgery and cannulation can be achieved—even though the postmortem intervals were longer than 15 min initially (pig A: 18.3 min; pig B: 22.8 min), we subsequently reduced it to below 10 min in pigs (pig C: 7.7 min; pig D: 8.7 min), and then determined that surgery and cannulation could also be achieved in less than 10 min in a human cadaver (9.9 min). In the final pig experiment (pig E) the postmortem interval was 13.8 min to approximate the upper limit of the perfusability window.

In pigs A and B, the postmortem interval was longer than the perfusability window, and the brains remained largely unfixed. In pigs C and D, postmortem interval was short, but histology revealed some areas of distorted morphology, especially in the white matter (Fig. 1A,E), which has a sparser capillary network than gray matter [21]. These areas showed decreased staining (pallor) at low power (Fig. 1A) and vacuolation at high power (Fig. 1E) in the white matter, similar to that observed for intramyelinic edema induced by toxicants [22] or disease [23]. Insufficient blood washout in these pigs was caused by imperfect placement of the cannula in the ascending aorta, and in both brains we observed erythrocytes in the brain capillaries (Fig. 1C). The correct cannula placement is made difficult by the fact that the ascending aorta is much shorter in pigs than it is in humans [24], and particular care needed to be taken to avoid positioning the cannula beyond the branching of the brachiocephalic artery.

**Fig. 1.**
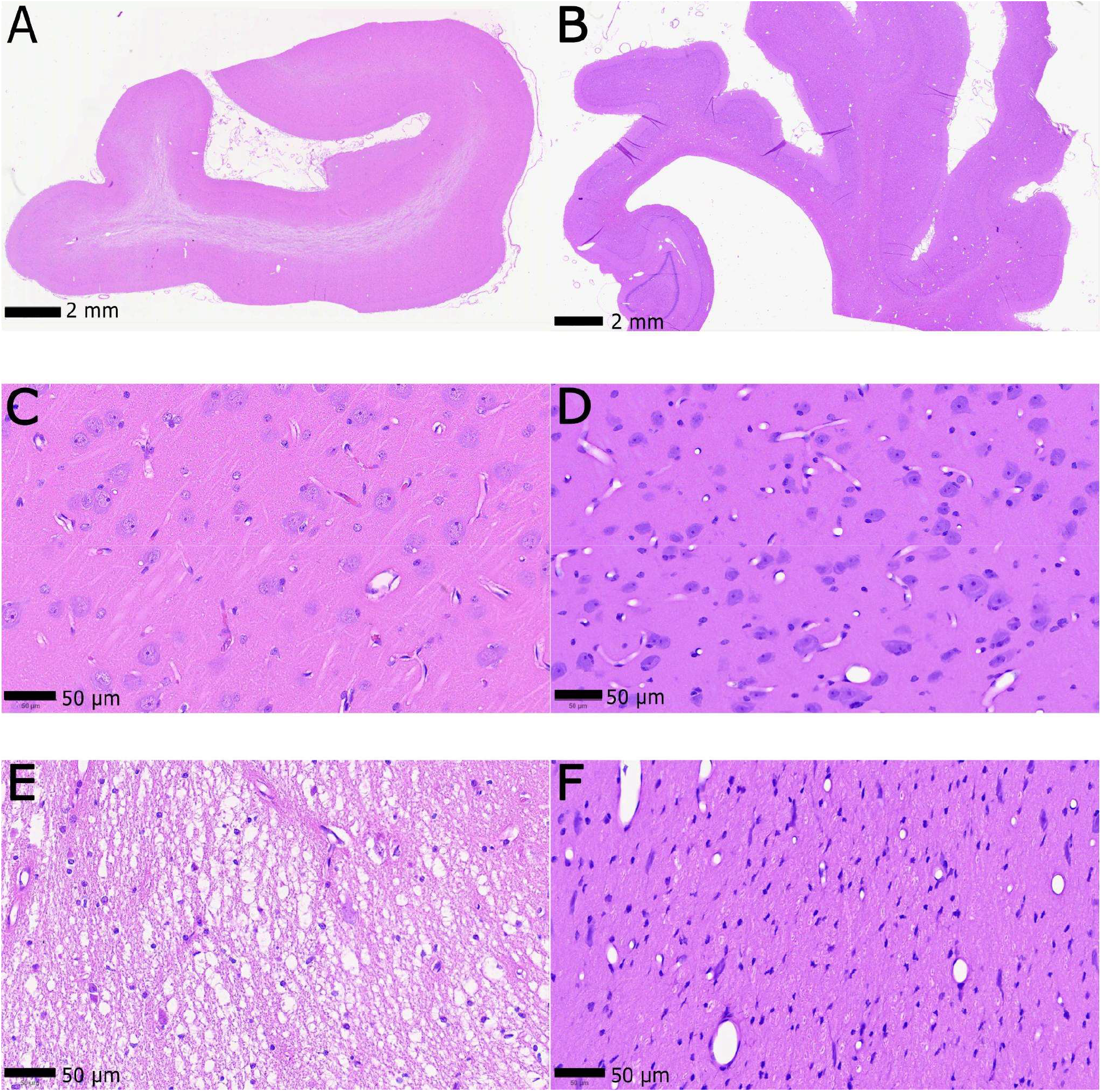
Hallmarks of imperfect ultrastructural preservation. A low power image of a section of neocortex from the right hemisphere of pig C reveals a visibly lighter region of hematoxylin and eosin staining (pallor; panel A); higher power images reveal erythrocytes in capillaries in the gray matter (C) and the appearance of vacuolation in the white matter (E). Such pallor and vacuolation are absent in the sections of neocortex from the right hemisphere from pig E (panel B), higher magnification shows empty capillaries in the gray matter (D) and no vacuolation in the white matter (F).

In the final experiment described here, pig E, the final concentration of ethylene glycol in the tissue was about 57 % w/w. On gross examination, the brain did not have any noticeable unfixed areas; and no distorted morphology was noticeable in hematoxylin and eosin staining in the sections of neocortex (Fig. 1B) nor other regions of the brain we analyzed (e.g, the cerebellum; Supplementary Materials).

The pig E brain shows excellent ultrastructural preservation (Fig. 2; Supplementary Materials): clearly visible cells with uniformly intact clear and compact membranes, and mitochondria that appear normal, with visible cristae and no examples of exploded and vacuolated (or ‘‘popcorned’’; [9,17]) mitochondria. No ‘‘dark’’ neurons [25] are present, indicating proper fixation and the lack of mechanical disruption to tissue. Several synapses can be clearly seen, with pre-synaptic vesicles and well defined, darkly stained post-synaptic densities. Slight loosening of the myelin can be observed around some processes, which however themselves were well defined, so connectomic analysis would still be possible. In volume electron microscopy (wafer-SEM and FIB-SEM; Fig. 2 and Supplementary Materials), it is easy to track the progression of any process through the stack.

**Fig. 2.**
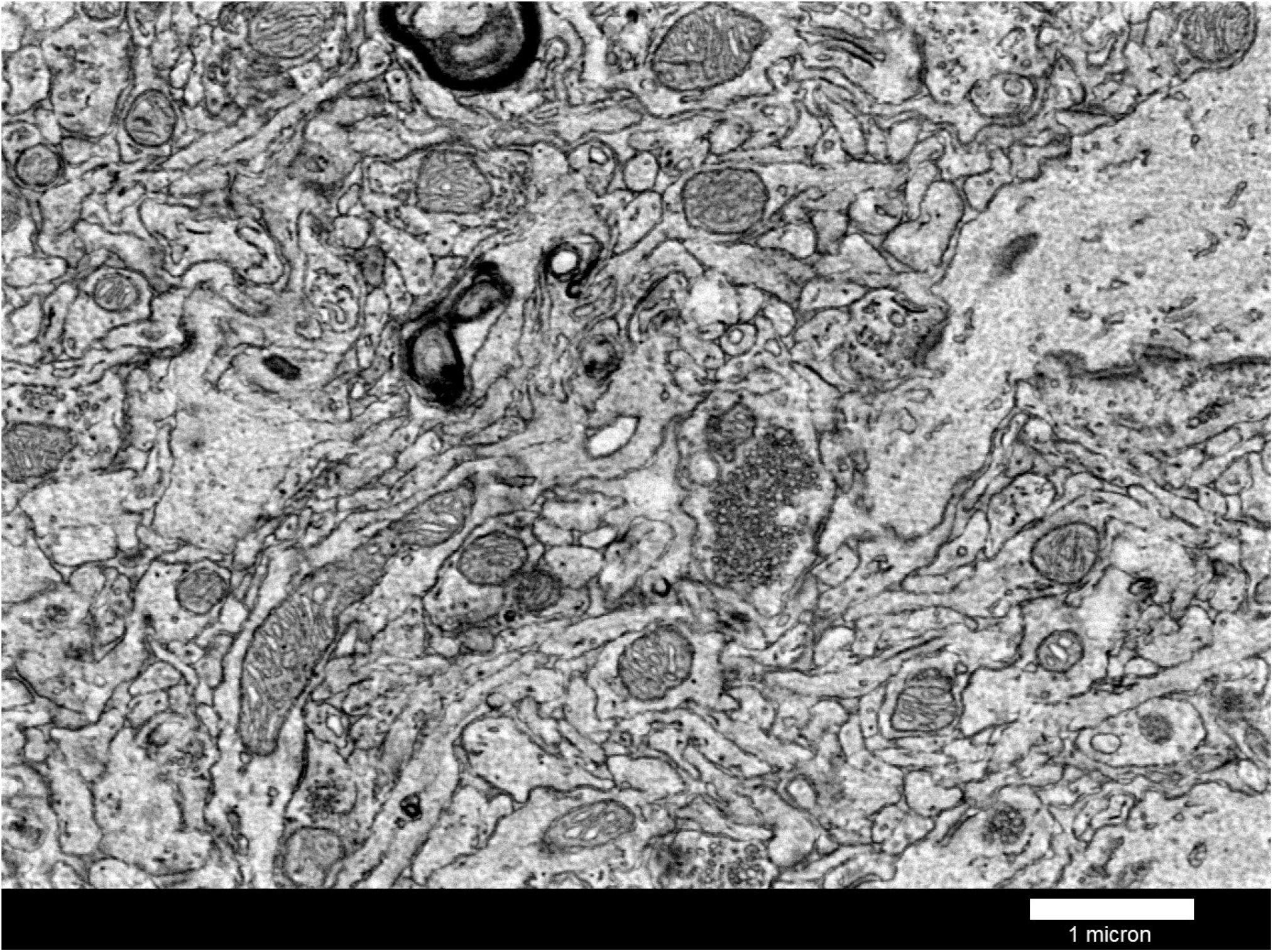
First section in a wafer-SEM stack of the neuropil from a randomly chosen area in the occipital lobe of pig E. All processes appear well preserved and traceable throughout the stack. The full stack is available in Supplemental Materials and was obtained by Ken Hayworth at Janelia Research Campus. Used with permission.

## Discussion

Our results show it is possible to introduce fixative for ultrastructural preservation, and then cryoprotectant, into large mammalian brains post-mortem.

We determined that there is but a short perfusability window in brain preservation. Although the size of this window in the human is not yet known, it is likely that it is of similar length, as the mechanisms responsible for the no-reflow phenomenon are expected to operate on a similar time scale. In our pig experiments, the perfusion needed to be initiated within about 15 min—less than 18 min, as shown by the failure of brain preservation in pig A, but possibly slightly longer than 14 min, as shown by the successful ultrastructural preservation in pig E. It seems that what matters here is the time during which the brain can be exposed to stagnant blood—other experiments [27] show that in pig brains obtained by rapid exsanguination, followed by flush-out of residual blood initiated within 10 min, perfusion could be restored even 240 min after cardiac arrest. Interestingly, exsanguination also appears to prolong the time after which cerebral recovery is possible following cardiac arrest in humans [28,29]. This time can also be prolonged by the use of barbiturates [28], which were used as part of end-of-life medication in our pig experiments, and are commonly used in human PAD [8].

Our experiments in the pig indicate that cannulation of the ascending aorta could be accomplished within 10 minutes of cardiac arrest, and thus within the perfusability window. Surgery and cannulation of human aortas could also be accomplished within this timeframe. The ultrastructure of the brain was not perfectly preserved in our initial pig experiments not because the surgery or cannulation took too long; rather, our results suggest it was because the blood washout was not perfect—that there was insufficient flow through the brain caused by imperfect placement of the cannula. Once we established this was the problem in some experiments, we determined that taking care to ensure that the cannula is placed correctly did not take significantly more time, adding at most 30 sec.

The histological analysis of imperfectly preserved brains show that distorted morphology is observed mostly in the regions of the brain that are less vascularized, namely, the white matter of the cerebral hemispheres. Poorer vascularization of the white matter [21] results in it receiving only ¼ of the blood flow received by gray matter—cerebral blood flow is 80 ml/min/100 g in gray matter and 20 ml/min/100 g in white matter [12]. It is likely that both during washout and fixation there is a similar proportion between the flow received by white and gray matter.

If the long-term connections, instantiated in the white matter [30], are not well-preserved, this precludes whole-brain reconstruction of neural connectivity. Because of the sheer number of these connections, it is unlikely that deciphering the molecular rules governing axon guidance and synaptic connectivity could make up for the missing information otherwise preserved in white matter ultrastructure. Although we believe that the vast majority of axons needs to be preserved, we do not dismiss the value of additional molecular information beyond structure. Reconstruction of connections is an error-prone process. The knowledge of what genes were expressed and proteins produced in neurons could provide barcoding data to improve traceability within a section and across sections [31] when, e.g., expansion microscopy is used as the imaging modality, although other modalities will probably be needed for human whole-brain connectomics [31,32]. Molecular information will also be helpful in deciphering the computational properties of individual neurons, if ultrastructure alone proves insufficient [33] and does not allow inference beyond basic functionality [34,35] on the path towards a whole human brain/body emulation [4–6]. We agree that structural—ultrastructural, to be precise—brain preservation (and especially, brain and body preservation) can provide a bridge to hypothetical restorative technologies in the future [36,37]. But contrary to the view expressed by McKenzie et al. [36], we find it implausible that immersion fixation or partial perfusion [7] would retain enough information to reach that goal. Still, we do agree that aldehyde fixation allows the maintenance of all the critical aspects: the shape of cell membranes and the location and structure (from primary to quaternary) of informational biomolecules (proteins and nucleic acids) [36].

Finally, we do not believe it is necessary to maintain the brains in solid state to prevent the deterioration of the brain ultrastructure and this molecular information. We demonstrated here and in our previous work [9] that it is possible to introduce cryoprotectant (such as ethylene glycol) to the brain of large mammals, through perfusion, without compromising brain ultrastructure, to concentrations allowing the elimination of crystallization during cooling and warming (metastable vitrification) [9,38]. However, storage below glass transition temperature—below -130°C for ethylene glycol:water mixtures [38]—would be very costly. Such mixtures allow not only for vitrification but also crystallization-free storage above the greatly depressed freezing temperature; there is a wide range of concentrations for which storage at around -35°C would be possible [39]. At a temperature this low, any non-enzymatic degradative process would be extremely slow, while processes catalyzed by enzymes would be stopped in fixed tissue, by direct deactivation of enzymes or immobilization of macromolecules [40]. For example, experimentally derived rate constants for non-enzymatic hydrolysis of peptide bonds [41], would result in only about 0.1% of bonds affected at -35°C in about 6 000 years. Cryoprotection of whole brains can also be of use for cutting sections of whole brains at cold temperatures [18,26].

## Limitations

We used a small number of pigs, and arbitrarily chose the sex of the animals—none were male. Although it is unlikely, this may decrease the generalizability of our findings. Pigs have similar body mass to humans, but smaller brains, about 100–160 g for adult pigs [42] vs. about 1000– 1600 g for adult humans [43,44]. Since the total cardiac output is similar, this means that a lower fraction of the total volume of the perfusate is received by the brain, reducing (about 10-fold) the probability of embolization per brain. More importantly, filtering effects of the carotid rete (rete mirabile)—a plexus of vessels that protects the pig cerebral circulation, absent in primates—is expected to further reduce this probability [45,46].

The rete may also buffer pressure surges—the presence of the rete is hypothesized to result in a more continuous blood flow in the brain [47,48]. However, we did not observe any such surges in our experiments, nor did we attempt to establish pulsatile flow. In fact, several elements of our perfusion circuit (filters, bubble traps) reduced small pressure amplitudes introduced by our pump. Although we found it unnecessary in our experiments, pulsatile flow—implemented, e.g., as in [27]—may provide advantages in brain preservation in humans.

Because we used young adult pigs, our experiments did not account for arterial pathologies (e.g., atherosclerotic plaque) and other co-morbidities that may be present in (mostly elderly) terminally ill humans and affect the timely initiation and lengthy continuation of perfusion. In particular, because cerebrospinal fluid is one of the main sources of fluid for anoxic cerebral edema and the volume of this fluid increases with age [49], elderly humans (and animals) may have a shorter perfusability window.

In our experiments, the end-of-life medication was administered intravenously, and the time from the drug administration to the cardiac arrest in our experiments was very short, less than 1 min. This is not always the case with medical aid-in-dying [50,8]. Prolonged agonal phase may result in poor perfusion of the organs [51], including the brain, and thus brain hypoxia, global or local [52], which may affect the length of the perfusability window.

## Conclusions

The protocol for post-mortem aldehyde-stabilized cryopreservation described in this paper is compatible with human brain preservation after physician-assisted death, provided that the blood washout is initiated less than about 14 min after cardiac arrest. We demonstrate, using light microscopy and volume electron microscopy, that this preservation protocol results in connectomically traceable whole brains that could be stored around -35°C, without degradation of ultrastructure or macromolecules, for thousands of years.

## Author Contributions

Conceptualization, A.S. and B.W.; Methodology, A.S., A.L., and B.W.; Investigation, A.S., A.L., and B.W.; Writing – Original Draft Preparation, B.W.; Writing – Review & Editing, A.S., A.L., and B.W.; Visualization, A.L. and B.W.; Funding Acquisition, A.S.

## Funding

This work was supported by the Biomedical Research and Longevity Society.

## Institutional Review Board Statement

Animal procedures were approved by the Institutional Animal Care and Use Committee at BioDevelopment Associates, Stanwood, WA (IACUC No.: 220601MP).

## Acknowledgments

We thank the employees of BioDevelopment Associates for their supervision of animal experiments, and Ken Hayworth (Janelia Research Campus) for generously providing wafer-SEM and FIB-SEM analysis of our brain samples.

## Conflicts of Interest

A.S., B.W., and A.L. are all employees of Nectome Inc., a for-profit research organization that offers high quality animal and human whole brain/body ultrastructural preservation.

## Supplementary Materials

High-resolution hematoxylin and eosin images of the sections of the neocortex of pig C and E, a section of the cerebellum of pig E, and wafer-SEM and FIB-SEM image stack for samples taken from the occipital cortex of pig E are available at https://github.com/boryswrobel-nectome/SongEtal2026-SupplementaryMaterial/

## Notes

https://github.com/boryswrobel-nectome/SongEtal2026-SupplementaryMaterial/

## References

1. Yang, W.; Yuste, R. Brain Maps at the Nanoscale. Nat Biotechnol 2019, 37, 378–380, doi:10.1038/s41587-019-0078-2.

2. Von Bartheld, C.S.; Bahney, J.; Herculano-Houzel, S. The Search for True Numbers of Neurons and Glial Cells in the Human Brain: A Review of 150 Years of Cell Counting. J of Comparative Neurology 2016, 524, 3865–3895, doi:10.1002/cne.24040.

3. Goriely, A. Eighty-Six Billion and Counting: Do We Know the Number of Neurons in the Human Brain? Brain 2025, 148, 689–691, doi:10.1093/brain/awae390.

4. Collins, L.T. The Case for Emulating Insect Brains Using Anatomical “Wiring Diagrams” Equipped with Biophysical Models of Neuronal Activity. Biol Cybern 2019, 113, 465–474, doi:10.1007/s00422-019-00810-z.

5. Amunts, K.; Axer, M.; Banerjee, S.; Bitsch, L.; Bjaalie, J.G.; Brauner, P.; Brovelli, A.; Calarco, N.; Carrere, M.; Caspers, S.; et al. The Coming Decade of Digital Brain Research: A Vision for Neuroscience at the Intersection of Technology and Computing. Imaging Neurosci 2024, 2, 1–35, doi:10.1162/imag_a_00137.

6. Linssen, C.; Koene, R. Functional Tests Guide Complex “Fidelity” Tradeoffs in Whole-Brain Emulation. J. Eth. Emerg. Tech. 2025, 35, 1–14, doi:10.55613/jeet.v35i1.152.

7. Garrood, M.; Keberle, A.; Sowa, A.; Janssen, W.; Thorn, E.; De Sanctis, C.; Farrell, K.; Crary, J.; McKenzie, A. Evaluating Ultrastructural Preservation Quality in Banked Brain Tissue. Free Neuropathology 2025, 1–13, doi:10.17879/FREENEUROPATHOLOGY-2025-6763.

8. Hoffman, D.N.; Strand, G.R. Clinical Practice and Pharmacology Decisions of Medical Aid-in-Dying Providers in the United States. BMJ Support Palliat Care 2025, spcare-2025-005397, doi:10.1136/spcare-2025-005397.

9. McIntyre, R.L.; Fahy, G.M. Aldehyde-Stabilized Cryopreservation. Cryobiology 2015, 71, 448–458, doi:10.1016/j.cryobiol.2015.09.003.

10. Lunney, J.K.; Van Goor, A.; Walker, K.E.; Hailstock, T.; Franklin, J.; Dai, C. Importance of the Pig as a Human Biomedical Model. Sci. Transl. Med. 2021, 13, eabd5758, doi:10.1126/scitranslmed.abd5758.

11. Patel, N.; Edwards, J.; Abdou, H.; Stonko, D.P.; Treffalls, R.N.; Elansary, N.N.; Ptak, T.; Morrison, J.J. Characterization of Cerebral Blood Flow during Open Cardiac Massage in Swine: Effect of Volume Status. Front. Physiol. 2022, 13, 988833, doi:10.3389/fphys.2022.988833.

12. Harrigan, M.R.; Leonardo, J.; Gibbons, K.J.; Guterman, L.R.; Hopkins, L.N. CT Perfusion Cerebral Blood Flow Imaging in Neurological Critical Care. NCC 2005, 2, 352–366, doi:10.1385/NCC:2:3:352.

13. Castaño, C.; Melià-Sorolla, M.; García-Serran, A.; DeGregorio-Rocasolano, N.; García-Sort, M.R.; Hernandez-Pérez, M.; Valls-Carbó, A.; Pino, O.; Grífols, J.; Iruela-Sánchez, A.; et al. Establishment of a Reproducible and Minimally Invasive Ischemic Stroke Model in Swine. JCI Insight 2023, 8, e163398, doi:10.1172/jci.insight.163398.

14. Hu, J.; Nan, D.; Lu, Y.; Niu, Z.; Ren, Y.; Qu, X.; Huang, Y.; Jin, H. Microcirculation No-Reflow Phenomenon after Acute Ischemic Stroke. Eur Neurol 2023, 86, 85–94, doi:10.1159/000528250.

15. Jia, M.; Jin, F.; Li, S.; Ren, C.; Ruchi, M.; Ding, Y.; Zhao, W.; Ji, X. No-reflow after Stroke Reperfusion Therapy: An Emerging Phenomenon to Be Explored. CNS Neurosci Ther 2024, 30, e14631, doi:10.1111/cns.14631.

16. Hallström, B.M.; Janke, A. Resolution among Major Placental Mammal Interordinal Relationships with Genome Data Imply That Speciation Influenced Their Earliest Radiations. BMC Evol Biol 2008, 8, 162, doi:10.1186/1471-2148-8-162.

17. Minassian, H.; Huang, S. Effect of Sodium Azide on the Ultrastructural Preservation of Tissues. Journal of Microscopy 1979, 117, 243–253, doi:10.1111/j.1365-2818.1979.tb01180.x.

18. Rosene, D.L.; Roy, N.J.; Davis, B.J. A Cryoprotection Method That Facilitates Cutting Frozen Sections of Whole Monkey Brains for Histological and Histochemical Processing without Freezing Artifact. J Histochem Cytochem. 1986, 34, 1301–1315, doi:10.1177/34.10.3745909.

19. Saija, A.; Princi, P.; Trombetta, D.; Lanza, M.; Pasquale, A.D. Changes in the Permeability of the Blood-Brain Barrier Following Sodium Dodecyl Sulphate Administration in the Rat: Exp Brain Res 1997, 115, 546– 551, doi:10.1007/PL00005725.

20. Knott, G.; Rosset, S.; Cantoni, M. Focussed Ion Beam Milling and Scanning Electron Microscopy of Brain Tissue. JoVE 2011, 2588, doi:10.3791/2588.

21. Wilhelm, I.; Nyúl-Tóth, Á.; Suciu, M.; Hermenean, A.; Krizbai, I.A. Heterogeneity of the Blood-Brain Barrier. Tissue Barriers 2016, 4, e1143544, doi:10.1080/21688370.2016.1143544.

22. Sills, R.C.; Johnson, G.A.; Anderson, R.J.; Johnson, C.L.; Staup, M.; Brown, D.L.; Churchill, S.R.; Kurtz, D.M.; Cushman, J.D.; Waidyanatha, S.; et al. Qualitative and Quantitative Neuropathology Approaches Using Magnetic Resonance Microscopy (Diffusion Tensor Imaging) and Stereology in a Hexachlorophene Model of Myelinopathy in Sprague-Dawley Rats. Toxicol Pathol 2020, 48, 965–980, doi:10.1177/0192623320968210.

23. Hayashi, M.; Ueda, M.; Hayashi, K.; Kawahara, E.; Azuma, S.; Suzuki, A.; Nakaya, Y.; Asano, R.; Sato, M.; Miura, T.; et al. Case Report: Clinically Mild Encephalitis/Encephalopathy with a Reversible Splenial Lesion: An Autopsy Case. Front. Neurol. 2024, 14, 1322302, doi:10.3389/fneur.2023.1322302.

24. Shah, A.; Goerlich, C.E.; Pasrija, C.; Hirsch, J.; Fisher, S.; Odonkor, P.; Strauss, E.; Ayares, D.; Mohiuddin, M.M.; Griffith, B.P. Anatomical Differences Between Human and Pig Hearts and Their Relevance for Cardiac Xenotransplantation Surgical Technique. JACC: Case Reports 2022, 4, 1049–1052, doi:10.1016/j.jaccas.2022.06.011.

25. Ahmadpour, S.; Behrad, A.; Vega, I.F. Dark Neurons: A Protective Mechanism or a Mode of Death. Journal of Medical Histology 2019, 3, 125–131, doi:10.21608/jmh.2020.40221.1081.

26. Kumarasami, R.; Verma, R.; Pandurangan, K.; Ramesh, J.J.; Pandidurai, S.; Savoia, S.; Jayakumar, J.; Bota, M.; Mitra, P.; Joseph, J.; et al. A Technology Platform for Standardized Cryoprotection and Freezing of Large-Volume Brain Tissues for High-Resolution Histology. Front. Neuroanat. 2023, 17, 1292655, doi:10.3389/fnana.2023.1292655.

27. Vrselja, Z.; Daniele, S.G.; Silbereis, J.; Talpo, F.; Morozov, Y.M.; Sousa, A.M.M.; Tanaka, B.S.; Skarica, M.; Pletikos, M.; Kaur, N.; et al. Restoration of Brain Circulation and Cellular Functions Hours Post-Mortem. Nature 2019, 568, 336–343, doi:10.1038/s41586-019-1099-1.

28. Breivik, H.; Safar, P.; Sands, P.; Fabritius, R.; Lind, B.; Lust, P.; Mullie, A.; Orr, M.; Renck, H.; Snyder, J.V. Clinical Feasibility Trials of Barbiturate Therapy after Cardiac Arrest: Critical Care Medicine 1978, 6, 228–244, doi:10.1097/00003246-197807000-00004.

29. Gilston, A. Complete Cerebral Recovery after Prolonged Circulatory Arrest: A Report of Two Cases. Intensive Care Med 1979, 5, 193–198, doi:10.1007/BF01683936.

30. Bullock, D.N.; Hayday, E.A.; Grier, M.D.; Tang, W.; Pestilli, F.; Heilbronner, S.R. A Taxonomy of the Brain’s White Matter: Twenty-One Major Tracts for the 21st Century. Cerebral Cortex 2022, 32, 4524– 4548, doi:10.1093/cercor/bhab500.

31. Collins, L.T.; Huffman, T.; Koene, R. Comparative Prospects of Imaging Methods for Whole-Brain Mammalian Connectomics. Cell Reports Methods 2025, 5, 100988, doi:10.1016/j.crmeth.2025.100988.

32. Wildenberg, G.; Boergens, K.; Nikitin, V.; Deriy, A.; De Carlo, F.; De Andrade, V.; Xiao, X.; Kasthuri, N. A Pipeline for a Primate Projectome: Mapping Every Individual Myelinated Axon across the Whole Brain 2023.

33. Zhang, Z.; Zhang, H.; Mihalas, S. Predicting Neural Activity from Connectome Embedding Spaces 2025.

34. Pospisil, D.A.; Aragon, M.J.; Dorkenwald, S.; Matsliah, A.; Sterling, A.R.; Schlegel, P.; Yu, S.; McKellar, C.E.; Costa, M.; Eichler, K.; et al. The Fly Connectome Reveals a Path to the Effectome. Nature 2024, 634, 201–209, doi:10.1038/s41586-024-07982-0.

35. Shiu, P.K.; Sterne, G.R.; Spiller, N.; Franconville, R.; Sandoval, A.; Zhou, J.; Simha, N.; Kang, C.H.; Yu, S.; Kim, J.S.; et al. A Drosophila Computational Brain Model Reveals Sensorimotor Processing. Nature 2024, 634, 210–219, doi:10.1038/s41586-024-07763-9.

36. McKenzie, A.T.; Zeleznikow-Johnston, A.; Sparks, J.S.; Nnadi, O.; Smart, J.; Wiley, K.; Cerullo, M.A.; De Wolf, A.; Minerva, F.; Risco, R.; et al. Structural Brain Preservation: A Potential Bridge to Future Medical Technologies. Front. Med. Technol. 2024, 6, 1400615, doi:10.3389/fmedt.2024.1400615.

37. McKenzie, A.T.; Wowk, B.; Arkhipov, A.; Wróbel, B.; Cheng, N.; Kendziorra, E.F. Biostasis: A Roadmap for Research in Preservation and Potential Revival of Humans. Brain Sciences 2024, 14, 942, doi:10.3390/brainsci14090942.

38. Faltus, M.; Bilavcik, A.; Zamecnik, J. Vitrification Ability of Combined and Single Cryoprotective Agents. Plants 2021, 10, 2392, doi:10.3390/plants10112392.

39. Cordray, D.R.; Kaplan, L.R.; Woyciesjes, P.M.; Kozak, T.F. Solid - Liquid Phase Diagram for Ethylene Glycol + Water. Fluid Phase Equilibria 1996, 117, 146–152, doi:10.1016/0378-3812(95)02947-8.

40. Migneault, I.; Dartiguenave, C.; Bertrand, M.J.; Waldron, K.C. Glutaraldehyde: Behavior in Aqueous Solution, Reaction with Proteins, and Application to Enzyme Crosslinking. BioTechniques 2004, 37, 790– 802, doi:10.2144/04375RV01.

41. Radzicka, A.; Wolfenden, R. Rates of Uncatalyzed Peptide Bond Hydrolysis in Neutral Solution and the Transition State Affinities of Proteases. J. Am. Chem. Soc. 1996, 118, 6105–6109, doi:10.1021/ja954077c.

42. Minervini, S.; Accogli, G.; Pirone, A.; Graïc, J.-M.; Cozzi, B.; Desantis, S. Brain Mass and Encephalization Quotients in the Domestic Industrial Pig (Sus Scrofa). PLoS ONE 2016, 11, e0157378, doi:10.1371/journal.pone.0157378.

43. Heymsfield, S.B.; Chirachariyavej, T.; Rhyu, I.J.; Roongpisuthipong, C.; Heo, M.; Pietrobelli, A. Differences between Brain Mass and Body Weight Scaling to Height: Potential Mechanism of Reduced Mass-Specific Resting Energy Expenditure of Taller Adults. Journal of Applied Physiology 2009, 106, 40– 48, doi:10.1152/japplphysiol.91123.2008.

44. Kandel, J.; Pokharel, D. Mean Brain Weight among Autopsy Cases at the Department of Forensic Medicine of a Tertiary Care Centre: A Descriptive Cross-Sectional Study. J Nepal Med Assoc 2022, 60, 274– 277, doi:10.31729/jnma.7162.

45. Haines, D.E.; Stewart, M.T.; Ahlberg, S.; Barka, N.D.; Condie, C.; Fiedler, G.R.; Kirchhof, N.A.; Halimi, F.; Deneke, T. Microembolism and Catheter Ablation I: A Comparison of Irrigated Radiofrequency and Multielectrode-Phased Radiofrequency Catheter Ablation of Pulmonary Vein Ostia. Circ: Arrhythmia and Electrophysiology 2013, 6, 16–22, doi:10.1161/CIRCEP.111.973453.

46. Kim, W.J.; Samarage, H.M.; Zarrin, D.; Goel, K.; Wang, A.C.; Johnson, J.; Nael, K.; Colby, G.P. Endovascular Transmural Access to Carotid Artery Perivascular Tissues: Safety Assessment of a Novel Technique. J NeuroIntervent Surg 2023, 15, 1007–1013, doi:10.1136/jnis-2022-019583.

47. Lluch, S.; Dieguez, G.; Garcia, A.L.; Gomez, B. Rete Mirabile of Goat: Its Flow-Damping Effect on Cerebral Circulation. American Journal of Physiology-Regulatory, Integrative and Comparative Physiology 1985, 249, R482–R489, doi:10.1152/ajpregu.1985.249.4.R482.

48. Reinert, M.; Brekenfeld, C.; Taussky, P.; Andres, R.; Barth, A.; Seiler, R.W. Cerebral Revascularization Model in a Swine. In New Trends of Surgery for Stroke and its Perioperative Management; Yonekawa, Y., Keller, E., Sakurai, Y., Tsukahara, T., Eds.; Acta Neurochirurgica Supplements; Springer-Verlag: Vienna, 2005; Vol. 94, pp. 153–157 ISBN 978-3-211-24338-1.

49. Du, T.; Mestre, H.; Kress, B.T.; Liu, G.; Sweeney, A.M.; Samson, A.J.; Rasmussen, M.K.; Mortensen, K.N.; Bork, P.A.R.; Peng, W.; et al. Cerebrospinal Fluid Is a Significant Fluid Source for Anoxic Cerebral Oedema. Brain 2022, 145, 787–797, doi:10.1093/brain/awab293.

50. Worthington, A.; Finlay, I.; Regnard, C. Efficacy and Safety of Drugs Used for ‘Assisted Dying. ’ British Medical Bulletin 2022, 142, 15–22, doi:10.1093/bmb/ldac009.

51. Silva E Silva, V.; Silva, A.R.; Rochon, A.; Lotherington, K.; Hornby, L.; Wind, T.; Bollen, J.; Wilson, L.C.; Sarti, A.J.; Dhanani, S. Organ Donation Following Medical Assistance in Dying, Part II: A Scoping Review of Existing Processes and Procedures. JBI Evidence Synthesis 2024, 22, 195–233, doi:10.11124/JBIES-22-00140.

52. Selkoe, D.J.; Myers, R.E. Neurologic and Cardiovascular Effects of Hypotension in the Monkey. Stroke 1979, 10, 147–157, doi:10.1161/01.STR.10.2.147.

